# Benchmarking beta-diversity measures and transfer functions for sedimentary ancient DNA

**DOI:** 10.1101/2025.08.15.670556

**Authors:** Tristan Cordier, François Keck, Anders Lanzén

## Abstract

Analyzing past ecosystems can improve our understanding of the mechanisms linking biodiversity with environmental changes. Sedimentary ancient DNA (*sed*aDNA) opens a window to past biodiversity, beyond the fossil record, that can be used to reconstruct ancient environments and ecosystems functions. To this end, modern biodiversity and environmental conditions are used to calibrate transfer functions, that are then applied to past biodiversity data to reconstruct environmental parameters. Doing this with *sed*aDNA can be challenging, because ancient DNA is often obtained in limited quantities and fragmented into smaller molecules. This leads to noisy datasets, with a low alpha diversity relative to modern DNA, patchy taxa detection patterns and/or skewed relative abundance profiles. How this affects beta-diversity measures, and the performance of transfer functions remain untested. Here we simulated ancient DNA reads counts matrices from synthetic and empirical datasets, and tested 464 combinations of counts transformations, beta-diversity indices, and ordinations methods, and assessed their performance in i) separating the ecological signal from the noise introduced by DNA degradation and in ii) predicting ground-truth environmental conditions. Our results show that commonly used workflows in DNA-based community ecology studies are sensitive to the noise associated to ancient DNA signal. Instead, combinations of methods that include more recent ordination methods proved robust to ancient DNA noise and produced better transfer functions. Our study provides a framework for designing post-processing workflows that are better suited for *sed*aDNA studies.

## Introduction

Documenting past environments and associated biodiversity is essential to improve our understanding of the dynamics of natural ecosystems over geological timescales. Current environmental reconstructions in paleoceanographic or paleolimnological studies commonly rely on transfer functions calibrated on microfossil assemblages [1]. For instance, assemblages of planktic foraminifera have been used to reconstruct Atlantic sea-surface temperatures [2, 3], dinoflagellate cysts for Arctic sea ice [4], and diatoms for lake salinity [5], trophic status [6] and water quality [7]. All of these methodologies rely on the ecological niche concept [8], which defines a theoretical multi-dimensional space where the dimensions represent environmental conditions under which a species can survive and reproduce. In practice, transfer functions exploit this principle by assuming a consistent and quantifiable relationship between species assemblages and environmental variables. Thus, species composition becomes a proxy for reconstructing past environmental conditions, based on the idea that taxa respond predictably to environmental gradients through their realized niches. Current transfer functions methodologies are currently limited to taxa that fossilize and can be identified in the sediment record. This means that on only a fraction of an ecosystem’s biodiversity can be considered, because soft-bodied taxa are not preserved in the sediment.

High-throughput sequencing of sedimentary ancient DNA (*sed*aDNA) is a powerful tool to study long-term biodiversity changes beyond the fossil record, as it provides direct molecular evidence of past communities preserved in environmental archives such as lake or marine sediments [9–12]. Over the last decade, *sed*aDNA datasets have been obtained at centennial timescales in lakes [13–15] and at glacial/interglacial (i.e. 10-200,000 years ago, [16, 17]) and even million-year [18, 19] timescales in marine ecosystems. These *sed*aDNA datasets thus open the path to leverage entire biodiversity datasets to reconstruct ancient environmental conditions. However, interpreting *sed*aDNA data remains challenging due to several biological and technical limitations. Among these, whether ancient DNA molecules from different taxa are evenly preserved in the sediment over time is not yet understood or documented, although sediments seem to present favorable physicochemical features for long-term DNA preservation [20, 21]. Such an uneven DNA preservation would result in missing taxa or patchy detections in *sed*aDNA datasets (i.e lower alpha diversity due to false negatives), and/or in skewed relative abundances profiles across samples (i.e. noisy beta-diversity). How this affects ecological inferences from beta-diversity patterns analysis, or environmental reconstructions via transfer functions, remain untested.

When analyzing biodiversity data obtained from DNA-based methods (i.e. metabarcoding or shotgun metagenomics), one needs to process the DNA read counts matrices, i.e. the community composition, via successive steps prior beta-diversity analysis or transfer function calibration. First, DNA read counts need to be transformed (or normalized), to compensate for the compositional nature of the data [22–24]. Then, these transformed counts are used to calculate pairwise community dissimilarities using a beta-diversity index [25, 26]. Finally, this dissimilarity matrix is used as input for an ordination method (usually a Non-metric Multidimensional Scaling, NMDS or Principal Coordinate Analysis, PCoA) to visualize beta-diversity patterns in a two-dimensional space [26]. While some counts transformation, beta-diversity indices and ordination methods have been developed decades ago, some more recent methods have been proposed to improve beta-diversity analyses or extract new ecological signal from high-dimensional datasets (Tables S1-S3), such as the ones produced with high-throughput sequencing technologies. For instance, the robust centered log ratio count transformation overcomes the need for adding a pseudocount to the matrix as it is the case with standard center log ratio transformation [27], and the wrench count transformation implements an empirical Bayes normalization approach to adjust for non-constant variance across samples [28]. Some beta-diversity indices now account for the phylogenetic signal extracted from DNA sequences to quantify community dissimilarities (i.e. UniFrac, [29] and PINA, [30]) or account for the taxa putative interaction patterns rather than only presence/absence (i.e. TINA, [30]), capturing new ecological dimensions from DNA-based datasets (Table S2). Newer ordination methods for high-dimensional data embedding (Table S3), such as t-Distributed Stochastic Neighbor Embedding (tSNE, [31]) and Uniform Manifold Approximation and Projection (UMAP, [32]) have recently proved useful for the analysis of beta-diversity patterns from DNA-based datasets [33, 34]. How the choice of the method for each of these of processing steps affect the results of a beta-diversity analysis has received substantial attention, but have mostly focused on either the effect of counts transformation [35–38], the effect of beta-diversity index [25] and the effect of the ordination method [33, 34]. No systematic benchmark of the effect of the multitude of possible combinations of these methods on the beta-diversity analysis has been carried out. Similarly, no benchmark currently exists on the effect of these processing steps on the performance of sample classification using supervised machine learning methods, or on the performance of transfer functions for environmental reconstructions with *sed*aDNA.

Here we simulated ancient DNA matrices from both synthetic and empirical community data to systematically benchmark 464 combinations of count transformation techniques, beta-diversity indices, and ordinations methods (Figure 1). We evaluated their performance in i) capturing the true ecological structure while compensating for technical noise introduced by the ancient DNA signal through beta-diversity patterns and ii) in producing accurate supervised machine learning models and transfer functions. Our goal is to identify the combinations of methods that maximize ecological signal and transfer function performance, even under the challenging constraints of *sed*aDNA data, such as possibly large rates of taxa dropout (or false negatives) due to DNA degradation over time.

**Figure 1:**
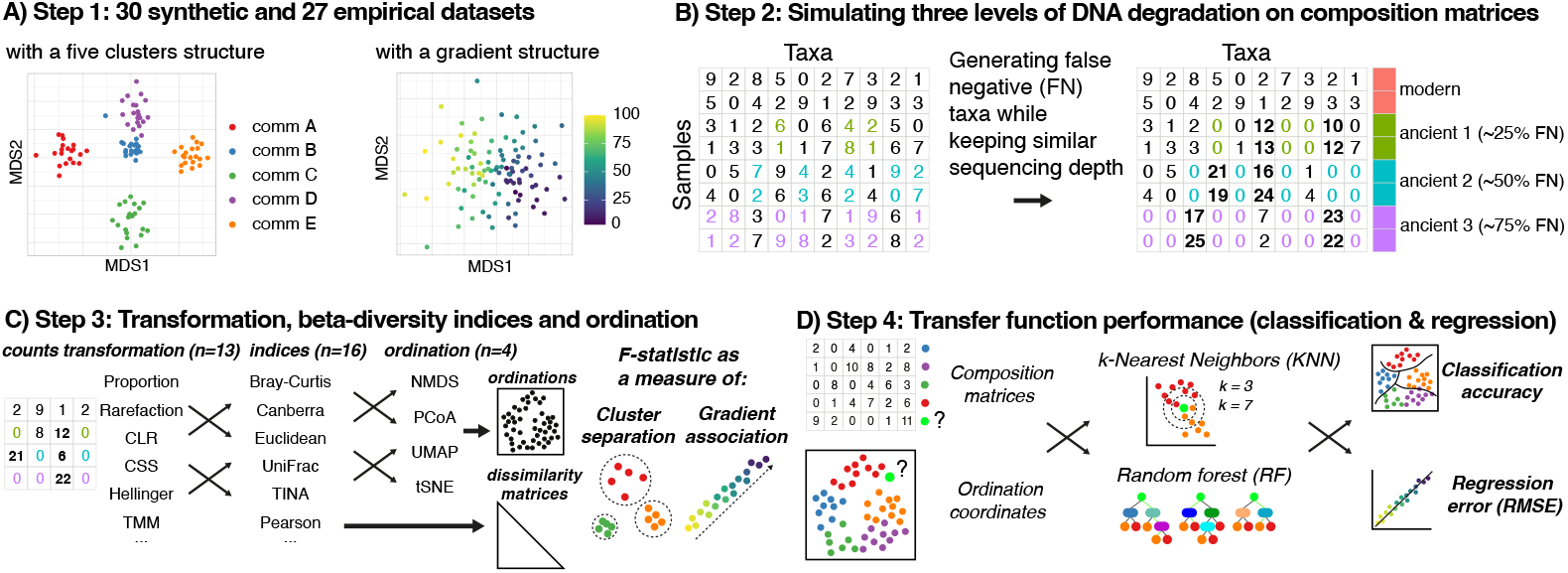
Overview of the approaches for data simulations and analysis. A) We produced 30 composition matrices using the splatter R package (synthetic datasets) and collected 27 environmental genomics datasets (empirical datasets, Table S4). B) Assuming that DNA degradation would generate false negatives taxa detections, we simulated three levels of false negative taxa, while keeping a similar sequencing depth (see methods for details). C) We produced 464 unique combinations of count transformation, beta-diversity indices and ordinations, and used the F-statistic of PERMANOVA models to measure their performance in separating clusters and find association with ecological gradients. D) We assessed the performance of these combinations of methods in producing accurate transfer functions, i.e. predicting correctly the cluster of origin and value on an ecological gradient for simulated ancient DNA samples.

## Material and methods

### Synthetic community datasets

We used the *splatter* R package v1.28.0 [39] to simulate 30 composition matrices comprising 100 samples and 400 taxa. Simulations were run both in a cluster (for community typing, n = 5 clusters) and in a path (for ecological gradient) setting, resulting in a total of 60 composition matrices (Figure 1A). For each simulation run, we used uniform distributions to generate a unique set of parameters that control the hierarchical probabilistic models from which the DNA read counts are drawn for populating composition matrices. Details of these simulations are available in the scripts deposited on the GitHub repository (https://github.com/trtcrd/Bench_sedaDNA), that allows reproducing the simulations as well as the downstream analysis.

### Empirical community datasets

We compiled 27 environmental genomics datasets (Table S4) of contrasted taxa (from bacteria to protists), in contrasting habitats (from pelagic to benthic and soil) and climates (from tropical to polar). We retrieved the composition matrices and associated metadata as they were produced in the respective studies. For datasets with a discrete variable (cluster analysis, n = 21), we balanced the number of samples per variable class to keep a fair comparison of classification performance. For datasets with a continuous variable (gradient analysis, n = 10), we kept all the original samples. To keep the computational needs of our benchmark reasonable, we reduced the number of samples to a maximum of 125 (Table S4). We also reduced the number of taxa to 1000-4000 taxa, by subtracting the same number of DNA reads across the entire composition matrices in order to keep the community structure. Finally, we kept only the samples with at least 20 taxa.

### Simulating ancient DNA signal

The expected signal of ancient DNA on community profiles is to have various levels of false negative taxa, i.e. taxa from which the DNA is not sequenced because it is not well preserved in the sediment record. Because the sequencing effort is a technical parameter, the DNA reads that are not dedicated to these false negative taxa would randomly cover the taxa from which DNA molecules are effectively present in the sample, and preferentially the most abundant ones. Under this rationale, we simulated three effect sizes of ancient DNA signal, by generating about 25, 50 and 75% false negatives taxa (allowing a small random noise around these values), randomly selected across the composition matrices (Figure 1B). This was done evenly across each variable class for community clusters or randomly along the ecological gradient, in order to have an even number of samples for each ancient DNA effect size (i.e. 25 % of samples at 0, 25, 50, 75% of false negatives). The reads subtracted from the false negatives were then allocated to a random selection of the 10% most abundant taxa of a given sample, to preserve the sequencing depth across the composition matrices.

### Statistics

We produced a total of 464 (synthetic datasets) and 396 (empirical datasets) combinations of count transformation techniques, beta-diversity indices and ordinations methods (Figure 1C, Table S1-3). For synthetic datasets, we mapped real plankton sequences obtained from a previous study [40] to the simulated taxa, using the same ranked relative abundances (i.e. most abundant plankton sequence from the study was mapped to the most abundant taxa in each synthetic dataset), allowing us to test combinations that include sequence-centered transformation techniques and beta-diversity indices (Table S2). We used the F-statistic of permutational analysis of variance (PERMANOVA) models as a measure of performance in cluster separation or gradient association, using the *adonis2* function of the *vegan* R package v2.6-6.1 [41]. This was measured from both samples pairwise dissimilarity matrices and from an Euclidean distance matrix extracted from the samples coordinates on the ordination’s embeddings. We also measured the changes in betadispersion, i.e. the multivariate variance, as a function of increasing ancient DNA effect size (from modern DNA to ancient DNA with 75% of false negative taxa) using the F-statistic output of the *betadisper* function of the vegan R package.

Then, we measured the performance of two supervised machine learning algorithms, namely k-nearest neighbors (KNN, [42]) and random forest (RF, [43]), in correctly classifying the samples (classification accuracy) or regressing the ecological gradients (root mean squared error, RMSE) using cross validation on each dataset. Gradients were normalized from 0 to 1 to keep the RMSE comparable across studies. Models were trained either on composition matrices (using the transformed DNA read counts of all taxa as features) or on ordination embeddings (using the samples coordinates in the two-dimensional space as features). We trained and tested the models using a 60-40 train-test split combined with repeated 10-fold cross-validation (10 repetitions) using the *caret* R package v6.0-94 [44], applied separately to each “modern DNA” and “ancient DNA” dataset.

We then measured the performance of KNN and RF-based transfer functions approaches (calibrated on composition matrices or ordination embeddings) in predicting the cluster of origin (transfer accuracy) or ecological gradient values (transfer RMSE) of the ancient DNA samples of each dataset, based on a model trained only on the modern DNA samples of that dataset (Figure 1D). Finally, we aggregated the performance metrics of the top five transfer function approaches and used Tukey’s Honest Significant Difference to test for differences in performance between KNN and RF-based transfer function approaches calibrated in either composition matrices or ordinations embeddings. Plots were done using the *ggplot2* R package v3.5.1 [45], and arranged with the *patchwork* R package v1.2.0 [46].

## Results

### Detecting clusters and gradients in distance matrices

Our results based on synthetic datasets show that the combinations of count transformation and beta-diversity indices that worked well for modern DNA corresponded, in general, to the ones that worked well for ancient DNA datasets (Figure 2A, 2C, Table S5-6). However, the combinations that included the TINA index worked particularly well for detecting clusters in modern DNA, while they performed poorly for ancient DNA. For both clusters and gradients, the Chi-Square transformation combined with the Pearson dissimilarity index (i.e. 1 – Pearson correlation between pairs of samples) resulted in the highest F-statistic across the 30 ancient DNA datasets. This was followed by several combinations of methods that included the Horn (i.e. 1 minus the Morisita-Horn index) or the Pearson index, that also worked well for modern DNA datasets. Interestingly, for empirical datasets, the two combinations of methods that included the TINA index resulted in the highest F-statistic for both modern and ancient DNA and for both cluster separation and gradient association, followed by multiple combinations of methods that included either the Pearson or Horn indices (Figure 2B, 2D, Table S7-8). For both synthetic and empirical datasets, the standard combinations of methods that are commonly used by environmental microbial ecologists (i.e. proportion and Bray-Curtis) or by human gut microbial ecologists (i.e. centered log ratio and Euclidean distance) were never among the best performing approaches for detecting clusters and gradients.

**Figure 2:**
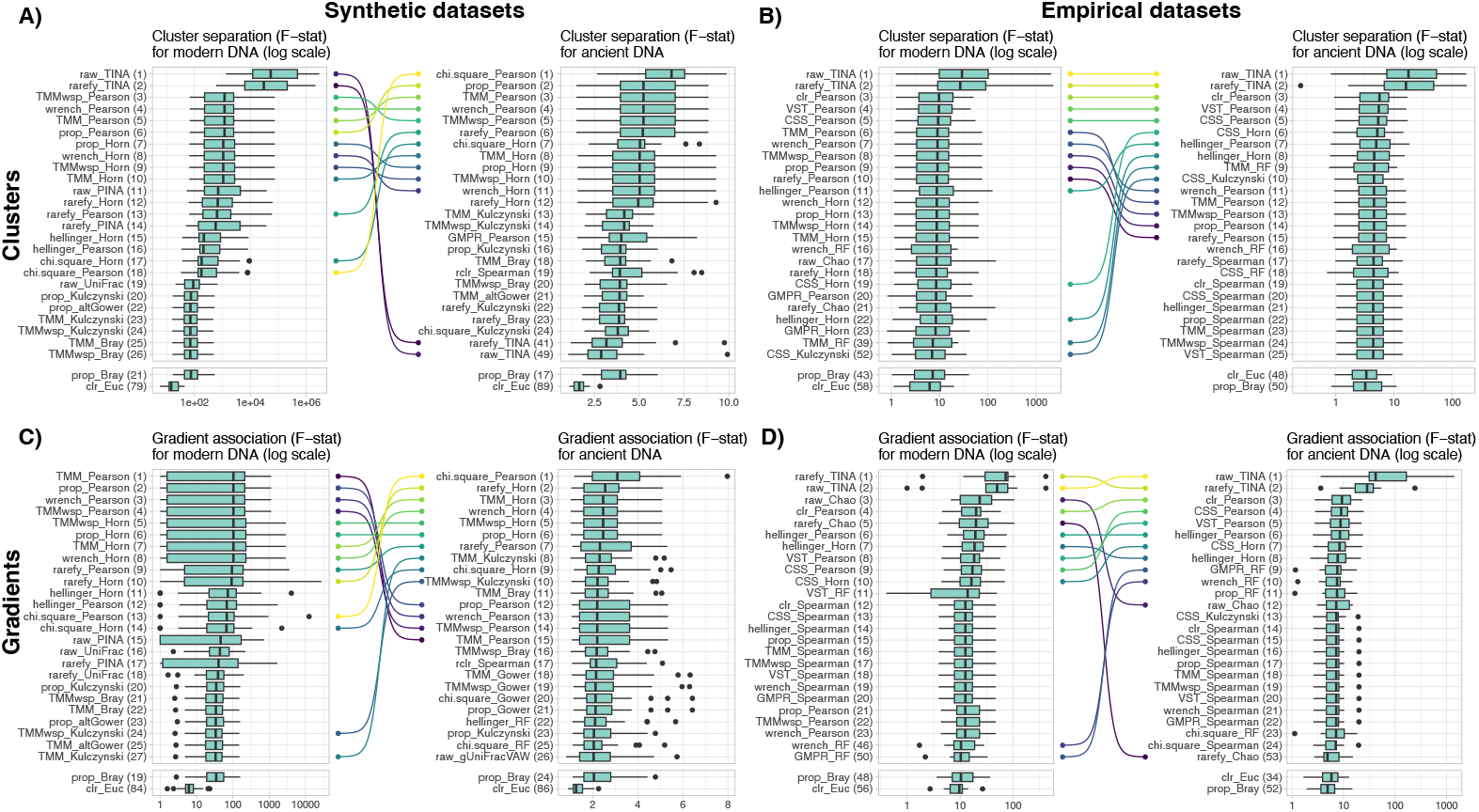
Performance of combinations of DNA read counts transformation and beta-diversity indices for cluster separation in A) synthetic datasets and B) empirical datasets, as well as of gradient association in C) synthetic datasets and D) empirical datasets. Performance was measured with the F-statistic of PERMANOVA models (higher is better). The left panels show the F-statistic distributions for modern DNA datasets, whereas the right panels show the F-statistic distributions for simulated ancient DNA datasets. Bumps charts connect identical combinations of methods between modern and ancient DNA datasets that are among the 10 best approaches. Numbers between brackets indicate the ranked position of the combinations of count transformation and beta-diversity indices based on the median F-statistic value.

### Detecting clusters and gradients in ordinations

Our results show that the combinations of count transformation, beta-diversity indices and ordinations performed differentially in detecting clusters and gradients in modern and ancient DNA datasets (Figure 3A-D, Table S9-12). The combinations that performed best for ancient DNA were less performant to detect clusters and gradients in modern DNA datasets, for both synthetic and empirical datasets. Interestingly, almost all combinations that performed well in detecting clusters and gradients in both modern and ancient DNA datasets included the UMAP ordination method. However, both UMAP and tSNE were ranked high for detecting gradients in modern synthetic and empirical DNA datasets (Figure 3C-D). Combinations that included the Chi-Square and Hellinger count transformation techniques were often ranked high in both synthetic and empirical datasets. Similarly, combinations that included the Canberra beta-diversity index for detecting clusters, and the Kulczynski or Bray-Curtis indices for detecting gradients, were often ranked at the top for synthetic datasets, while it was the Spearman index that performed well for empirical datasets, and to a lesser extent the Kulczynski or Bray-Curtis indices. The standard combinations of methods for beta-diversity analysis in microbial ecology (i.e. proportion, Bray-Curtis and NMDS or centered log ratio, Euclidean and PCoA) were ranked much lower for both synthetic and empirical and for both modern and ancient DNA datasets.

**Figure 3:**
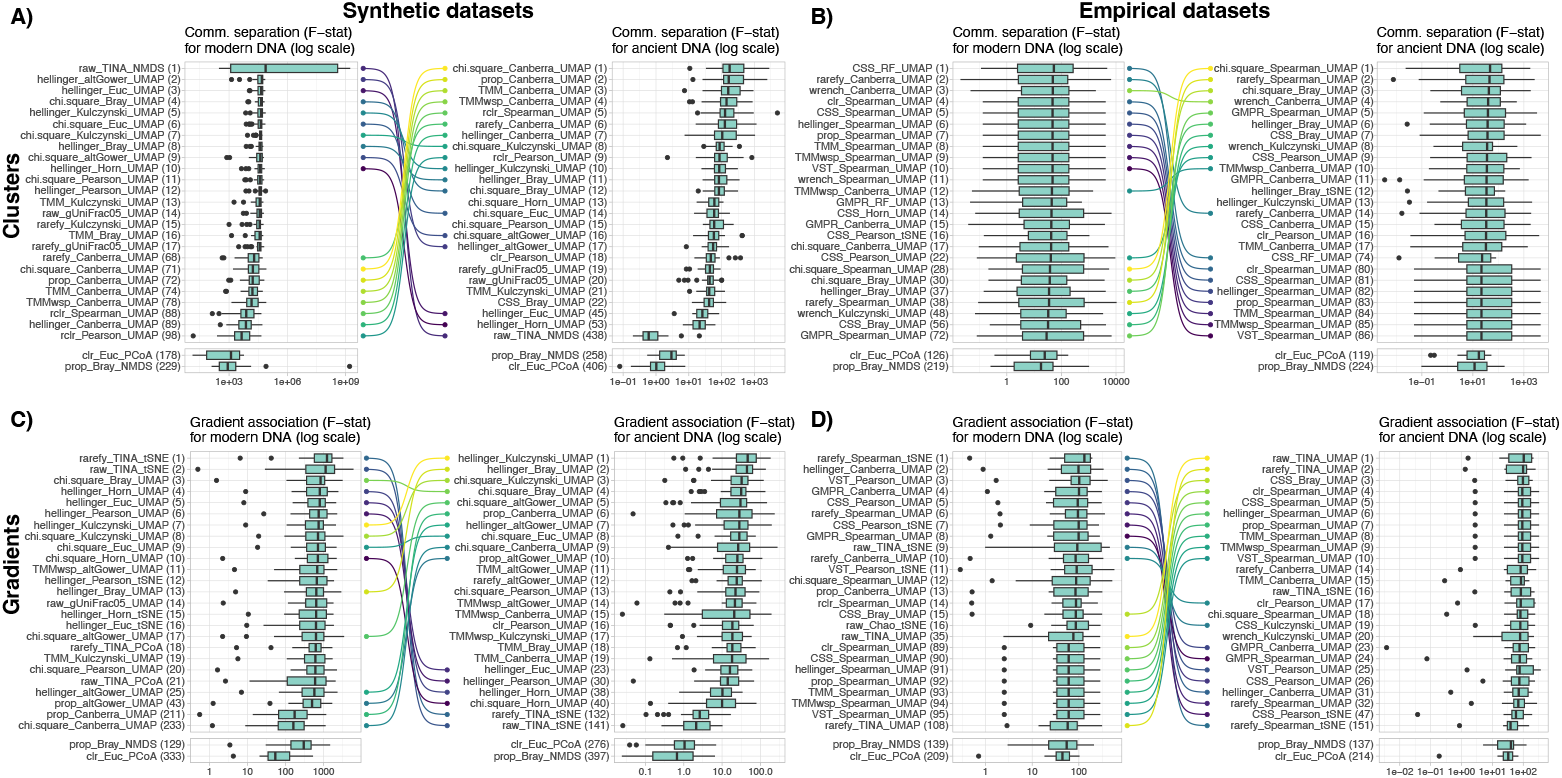
Performance of combinations of DNA read counts transformation, beta-diversity indices and ordinations for cluster separation in A) synthetic and B) empirical datasets, and for gradient association in C) synthetic and D) empirical datasets. Performance was measured with the F-statistic of PERMANOVA models (higher is better). The left panels show the F-statistic distributions for modern DNA datasets, whereas the right panels show the F-statistic distributions for simulated ancient DNA datasets. F-statistics were calculated using the Euclidean distances between pairs of samples in the ordinations as input of PERMANOVA models. Bumps charts connect identical combinations of methods that are among the 10 best approaches. Numbers between brackets indicate the ranked position of the combinations of count transformation, beta-diversity index and ordination based on the median F-statistic value.

### Cross validation of supervised machine learning models and transfer functions performance

We measured the performance of KNN and RF supervised models to predict the cluster of origin (classification accuracy), or the ecological gradient value (RMSE), for each sample on both modern and ancient DNA datasets using repeated 10-fold cross validations. Models were trained either on composition matrices (using the transformed DNA read counts as features, Figure 4) or on ordination embeddings (using the samples coordinates in the two-dimensional space as features, Figure 5).

**Figure 4:**
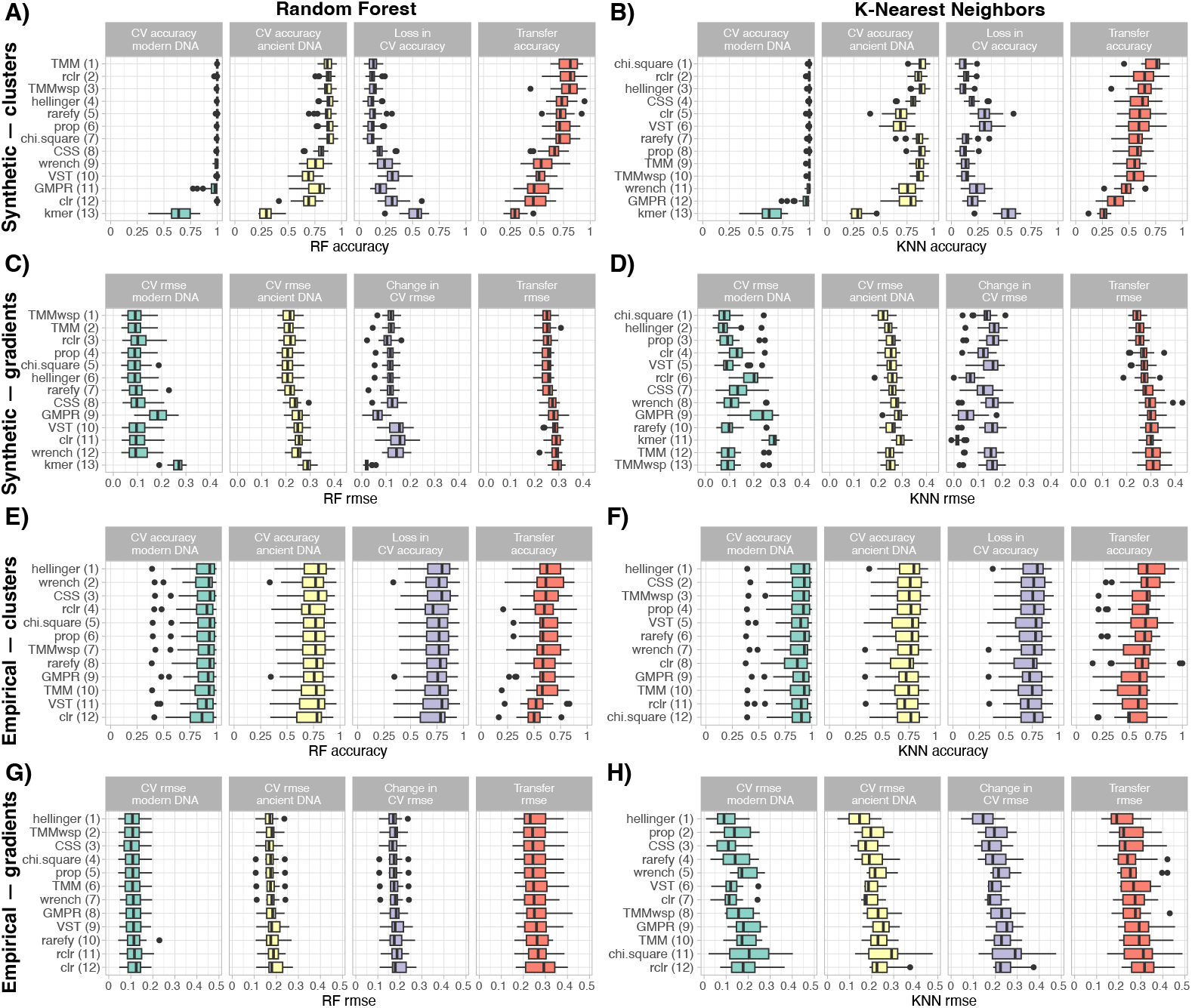
Performance of supervised models and transfer functions trained on composition matrices, i.e. using transformed DNA read counts as features, to infer the cluster of origin for synthetic (A-B) and empirical (E-F) datasets, and the ecological gradient value for synthetic (C-D) and empirical (G-H) datasets. We measured the performance of supervised models on both modern and ancient DNA datasets by cross validation (CV on two first panels, with a third panel showing the change in accuracy for clusters or in root mean square error for gradients between modern and ancient DNA). We finally measured the performance of transfer functions (the fourth panel, in red) trained only on modern samples to infer the cluster of origin and ecological gradient values on the ancient DNA samples. Numbers between brackets indicate the ranked performance of the count transformation techniques based on the median value of transfer accuracy-RMSE.

**Figure 5:**
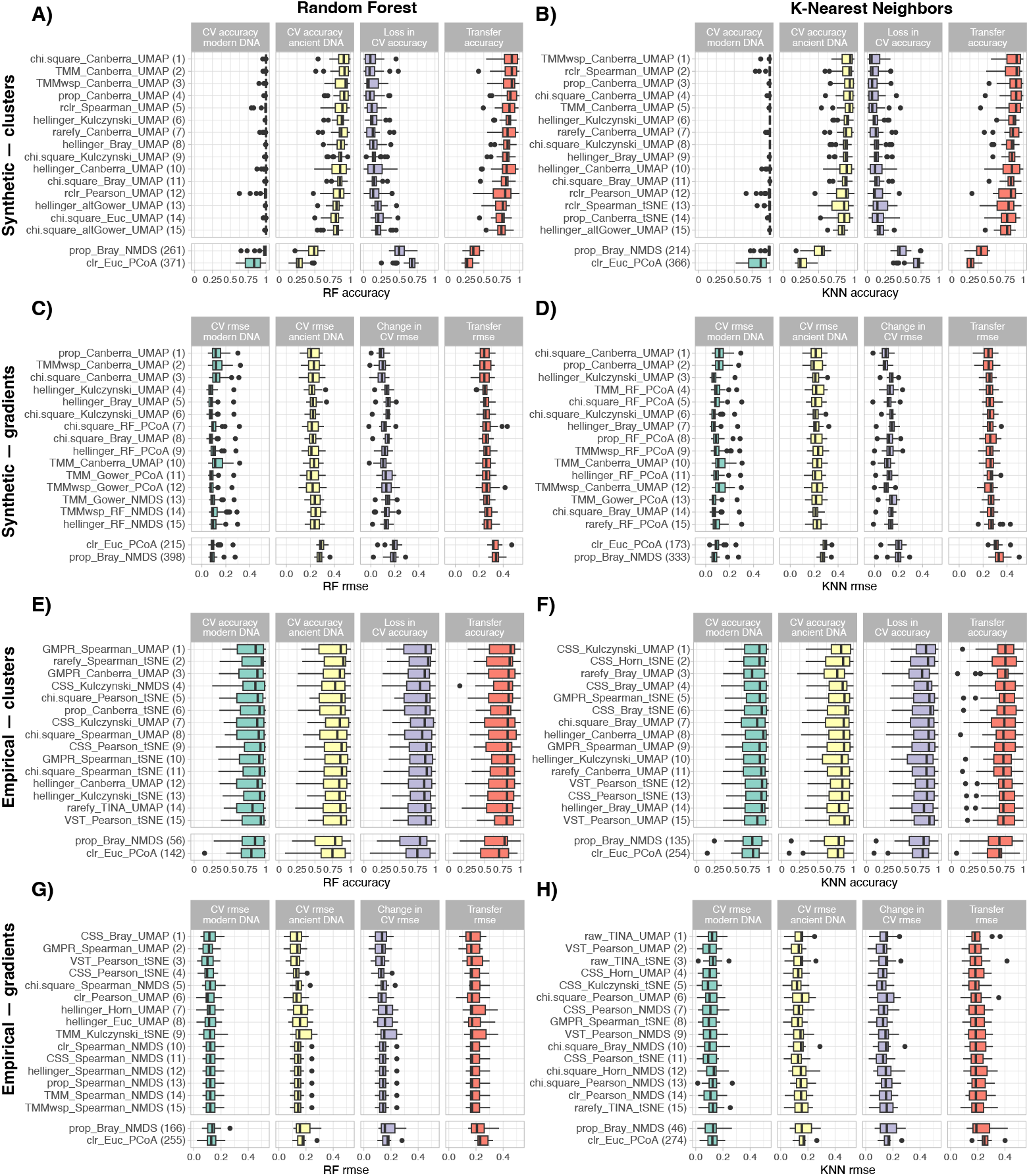
Performance of supervised models and transfer functions trained on ordinations embeddings, i.e. using samples coordinates on the ordination as features, to infer cluster of origin for synthetic (A-B) and empirical (E-F) datasets, and the ecological gradient value for synthetic (C-D) and empirical (G-H) datasets. We measured the performance of supervised models on both modern and ancient DNA datasets by cross validation (CV on two first panels, with a third panel showing the change in accuracy for clusters or in root mean square error for gradients between modern and ancient DNA). We finally measured the performance of transfer functions (the fourth panel, in red) trained only on modern samples to infer the cluster of origin and the ecological gradient values on the ancient DNA samples. Numbers between brackets indicate the ranked performance of the combinations of count transformation techniques, beta-diversity indices and ordinations methods based on the median value of transfer functions accuracy-RMSE.

For composition matrices, supervised models cross-validated on synthetic modern DNA datasets showed that count transformation techniques have only little effect on performance, except for the kmer transformation and to a lesser extent the GMPR transformation (Figure 4A-D, Table S13-14). Performance on empirical datasets was mostly similar across the different count transformations techniques (Figure 4E-H, Table S15-16). Supervised models cross-validated on synthetic ancient DNA datasets show that the kmer, clr, GMPR, VST and wrench count transformation techniques are less performant than others, especially for cluster inference (Figure 4A-D). Again, this was not the case for ancient DNA datasets obtained from empirical datasets, where the count transformation technique showed little to no effect (Figure 4E-F), although the Hellinger transformation gave better results for empirical gradients (Figure 4H). Finally, the count transformations techniques identified above also produced less performant transfer functions, especially for cluster inference in synthetic datasets (Figure 4A-D). The effect of count transformation techniques on transfer function performance on empirical datasets has only a limited effect (Figure 4E-H).

For ordination embeddings, supervised models cross-validated on both synthetic and empirical modern and ancient DNA datasets show that combinations of count transformation techniques, beta-diversity indices and, to a lesser extent, ordinations methods only slightly affect performance (Figure 5A-H, Table S17-20). The same was observed for the performance of transfer functions, although here the ordination method had a stronger effect on performance (Figure S2). For cluster inferences, the combinations of methods including UMAP, and to a lesser extent tSNE ordinations, typically performed the best. For gradient inferences, combinations of methods including UMAP or PCoA performed best on synthetic datasets (Figure 5C-D), whereas UMAP, tSNE or NMDS are usually the best for empirical datasets (Figure 5G-H). The performance of transfer functions produced using standard combinations of methods in microbial ecology (i.e. proportion, Bray-Curtis and NMDS and center log ratio, Euclidean and PCoA) is much lower. For synthetic datasets, the combination of proportion, Bray-Curtis and NMDS was comparatively much better for clusters inference, whereas center log ratio, Euclidean and PCoA was better for gradients inference (Figure 5A-D). For empirical datasets, the performance of these combinations was similar for both cluster and gradient inference (Figure 5E-H).

Analyzing the five best composition and ordination-based transfer function approaches as a function of DNA effect size showed that the performance tends to drop as the ancient DNA effect sizes increases (Figure 6A-H). However, the performance of transfer functions calibrated on ordinations obtained from empirical datasets with the KNN algorithm is mostly similar at all ancient DNA effect sizes (Figure 6F, Figure 6H). We aggregated transfer accuracy and RMSE from these top five method combinations to compare the performance of composition-based and ordination-based transfer functions calibrated with either the random forest or k-nearest neighbors algorithms across the different levels of ancient DNA effect size. The best transfer function approach was not the same as the ancient DNA effect size increases (Figure 7). For cluster and gradient inferences on both synthetic and empirical datasets, the performance of ordination-based transfer functions (using either RF or KNN) is usually better, especially as the ancient DNA effect size increases (Figure 7A, Figure 7C).

**Figure 6:**
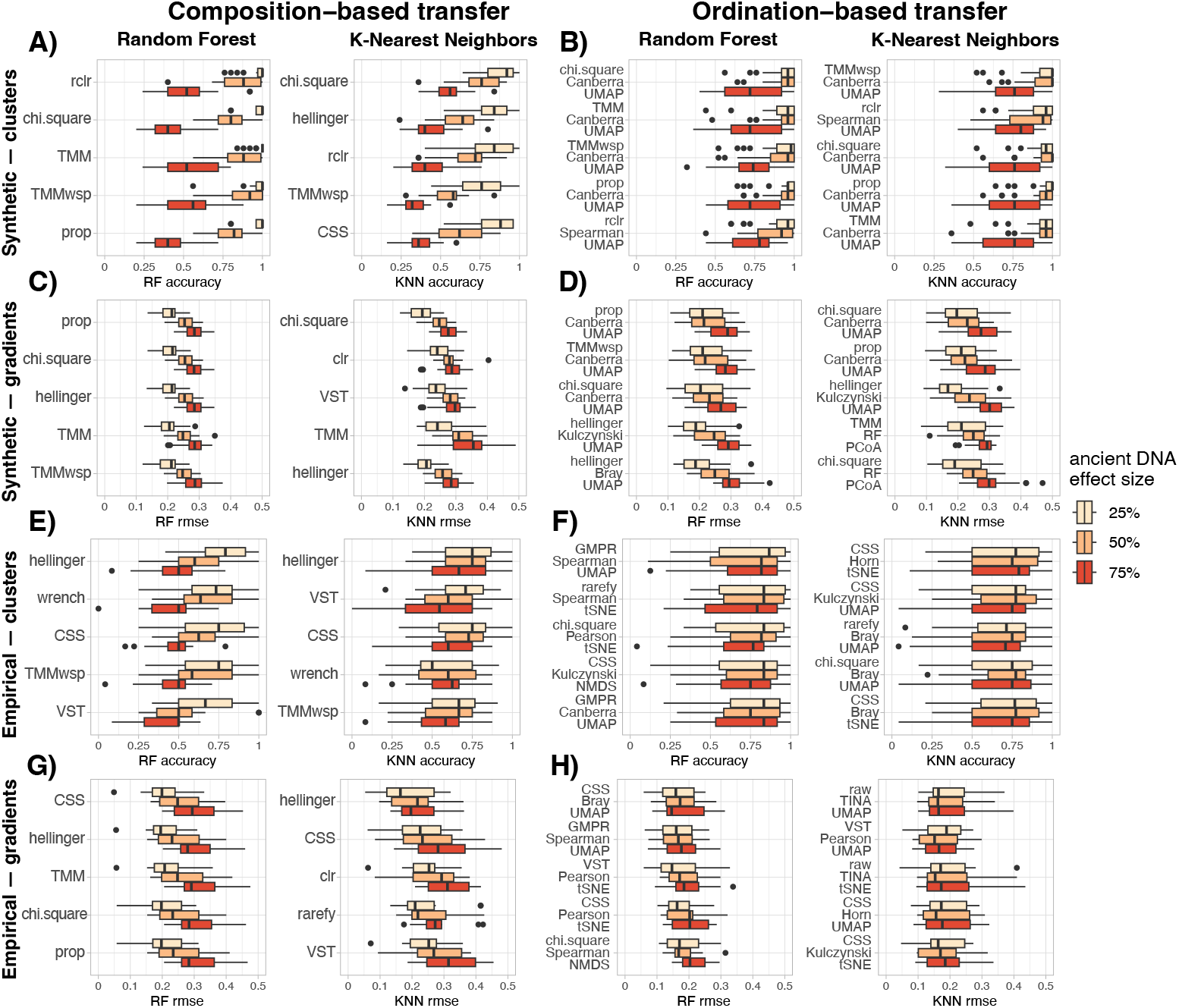
Performance of the top five composition-based and ordination-based transfer functions approaches in inferring cluster of origin for synthetic (A-B) and empirical (E-F) datasets, and the ecological gradient value for synthetic (C-D) and empirical (G-H) datasets, as a function of ancient DNA effect size.

**Figure 7:**
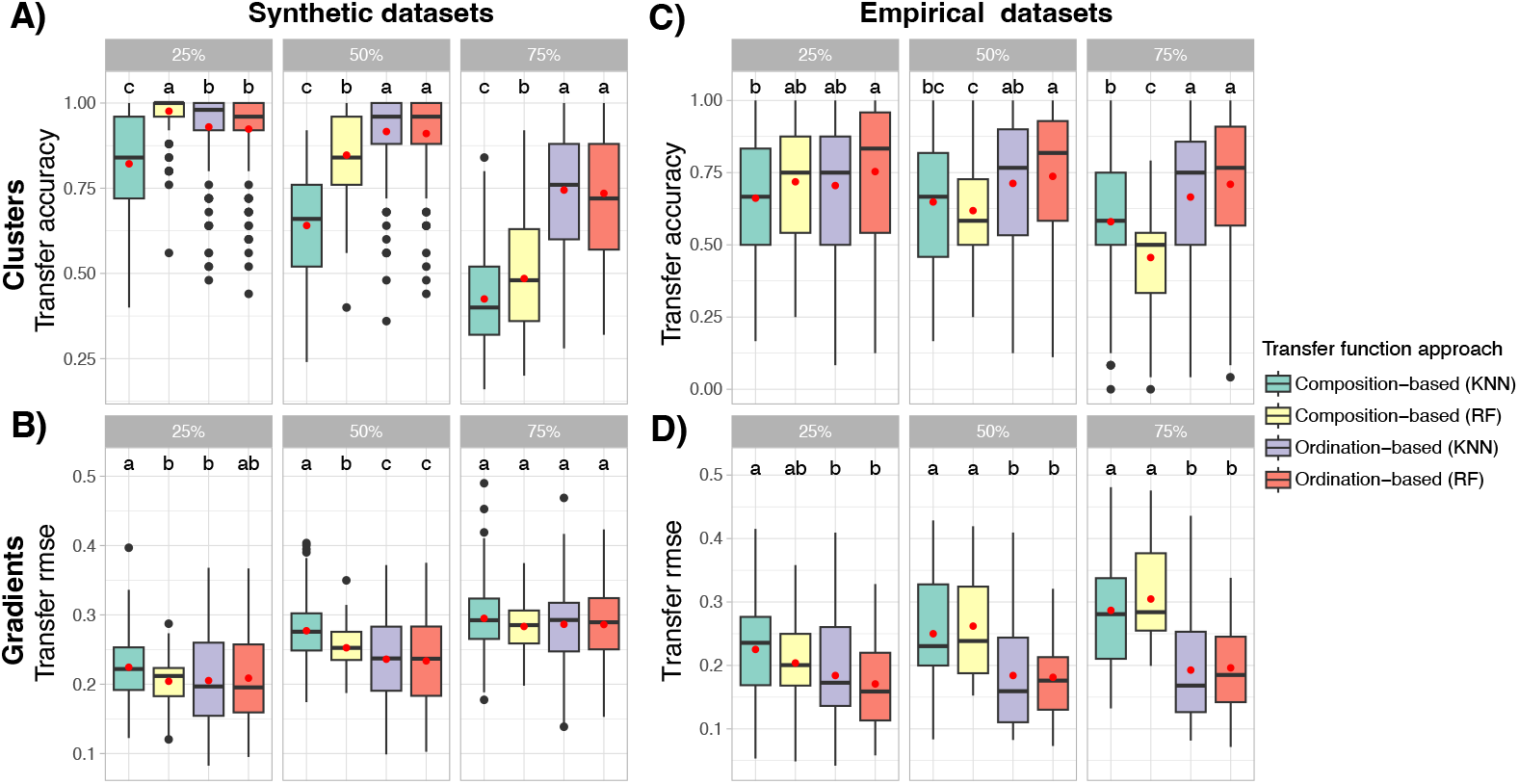
Aggregated performance metrics of the top five composition-based and ordination-based transfer functions approaches in inferring cluster of origin for synthetic (A) and empirical (C) datasets (transfer accuracy, higher is better), and the ecological gradient value for synthetic (B) and empirical (D) datasets (transfer RMSE, lower is better), as a function of ancient DNA effect size. We aggregated the performance metrics obtained from either random forest (RF) or the k-nearest neighbors (KNN) algorithms. We then compared the distributions of transfer functions accuracy-RMSE values using Tukey’s Honest Significant Difference to identify the best-performing transfer function approach at each level of ancient DNA effect size. Letters above the boxplots indicate groupings based on Tukey’s HSD: transfer function approaches that share the same letter(s) are not significantly different from each other (p > 0.05).

## Discussion

### Performance of beta-diversity measures from community dissimilarity matrices

Our benchmark of beta-diversity measures from community dissimilarity matrices shows that Pearson and Horn indices were among the best at detecting clusters and gradients in both modern and ancient DNA (Figure 2). Although the combination of Chi-Square and Pearson index was ranked first for ancient DNA in synthetic datasets, this was not the case for modern DNA datasets, where it ranked 18^th^ and 13^th^ for cluster and gradient, respectively. On empirical datasets, the approach combining Chi-Square and Pearson index was not among the top 25 combinations. Instead, combinations that included the TINA index were the best at detecting clusters on both modern and ancient DNA datasets, followed by combinations of methods that included the Person index (Figure 2). Our results are in line with previous work on modern DNA datasets, including the study that introduced the TINA beta-diversity index and compared its performance with commonly used indices [30], and a benchmark of beta-diversity indices based on simulated and empirical datasets that identified the Pearson index as particularly performant in detecting gradients [25]. Here, the TINA index did not perform well at detecting clusters in ancient DNA versions of the synthetic datasets, whereas it was included in the best combinations for empirical datasets. This difference may stem from the fact that our empirical datasets had two to ten times more taxa than synthetic datasets, which only comprised 400 taxa. Since the TINA index includes a step to infer putative taxa interactions as part of the community dissimilarity measure, having more taxa may help detecting interactions and thus contribute to better separate clusters. Thus, we produced synthetic datasets with 2000 taxa and rerun the benchmark on these larger matrices. We then observed that the combination of rarefaction with the TINA index was the best at detecting clusters (Figure S1). This supports the idea that the TINA index performs especially well in highly dimensional datasets, which is often the case in microbiome datasets. However, the TINA index includes the inference of an interaction network, which can be very computationally intensive. Since the Pearson and Horn indices were also identified as highly performant in detecting clusters and gradients, we recommend using these indices in highly dimensional datasets if computing power is limiting.

Finally, our results indicate that the most popular combinations of count transformation techniques and beta-diversity indices (i.e. taxa proportion and Bray-Curtis index, or centered log ratio and Euclidean distances) are never among the top ten approaches for beta-diversity measures and perform significantly worse than other combinations tested here. This is also in line with previous work [25, 30]. The common use of these approaches may come from historical practices [47] or from a theoretical standpoint [24, 48], and should be questioned based on the results presented here.

### Performance of beta-diversity measures and sample classification from community ordinations

Our results clearly show that combinations including UMAP, and to a lesser extent tSNE, almost always give the best results in detecting clusters and gradients in both modern and ancient DNA versions of all the tested datasets (Figure 3) as well as in sample classification (or regression) with cross-validated supervised models (Figure 5, Figure S2). These ordinations methods consistently outperform the commonly used NMDS and PCoA that are currently the standard approach for DNA-based community ecology studies (Figure 3, Figure S2). Our results show that the choice of the ordination method has a much stronger effect on beta-diversity measures and sample classification compared to count transformation and beta-diversity index (Figure S2). Combinations including UMAP and tSNE allowed extracting relevant ecological signals while compensating for technical noise due to ancient DNA signal introduced here, whereas combinations including NMDS and PCoA were sensitive to this noise. Indeed, NMDS ordinations were not separating the samples according to the DNA status (modern or ancient DNA), but were particularly prone to an increase in betadispersion (i.e. multivariate variance) as the ancient DNA effect size increases (Figure S2, Figure S3). PCoA ordinations tended to separate samples based on the DNA status, rather than cluster of origin (Figure S2, Figure S4). UMAP and tSNE were both able to extract the ecological signal while compensating for technical noise due to ancient DNA signal (Figure S2, Figure S5). These differences likely contribute to explain the variations in performance for classifying (or regressing) samples by cross-validated supervised models in both modern and ancient DNA datasets (Figure S2). These results are in line with previous work showing that UMAP proved superior to other ordinations methods in extracting relevant ecological signal in DNA-based microbiome datasets [33, 34]. Although not new to other disciplines utilizing high-throughput DNA sequencing technologies (e.g. for RNAseq analysis, [49]), UMAP and tSNE are currently underused in DNA-based community ecology studies. We therefore recommend these ordinations techniques in future studies, especially for ancient DNA data or in datasets with large variations in alpha diversity. Indeed, datasets with large variation in alpha diversity are prone to the horseshoe effect, where the samples are ordinated along a curve on the ordination space, because some samples may not share enough common taxa [50].

### Transfer functions based on ordination outperform the ones based on composition for sedaDNA datasets

The performance of transfer functions decreased as the ancient DNA effect size increased (Figure 6), as expected given the incremental addition of technical noise in the data. However, transfer functions calibrated on ordinations were still sufficiently performant with about 50% of false negative taxa in synthetic datasets (Figure 6B, 6D), and similarly performant across all ancient DNA effect sizes in empirical datasets (Figure 6F, 6H). Inspecting the aggregated performance metrics obtained from the top five combinations of methods for composition and ordination-based transfer functions confirmed that using ordinations for calibrating a transfer function gives better results, either with RF or KNN algorithms (Figure 7). The gap in performance between these two approaches was increasing as the ancient DNA effect size increased, supporting the idea that ordinations (especially UMAP and tSNE) can compensate for technical noise due to ancient DNA signal. We thus recommend using transfer functions that are calibrated on samples coordinates obtained from community ordinations for *sed*aDNA studies.

### Limitations and future work

Here, we simulated ancient DNA signal on both “modern” synthetic and empirical datasets. We based these simulations on the idea that *sed*aDNA samples may have “lost” some or many taxa over time, due to DNA degradation once buried in the sediment. These taxa thus cannot be detected in *sed*aDNA datasets and constitutes false negative detections that introduces noise in data analysis. Since the environmental conditions and processes affecting DNA taphonomy in the sediment are not yet fully understood (but see [20]), our simulations may only partially capture these factors and thus might not fully reproduce realistic *sed*aDNA noise patterns in our datasets. Furthermore, we did not account for possible false positives taxa detections, due to “tag-jumps” that can cause incorrect assignment of sequences to samples in metabarcoding studies [51, 52]. Lastly, we simulated ancient DNA noise to our datasets, but we did not change the community structure in any way. Thus, our simulated ancient DNA datasets do not account for possible changes in taxa ecology over time and their resulted assembly into potentially different communities relative to modern conditions. To overcome these limitations, future work would need to produce *sed*aDNA datasets alongside conventional proxy data that are used for paleoceanographic or paleolimnological reconstructions. Nevertheless, our study provides the first systematic benchmark of post-processing workflows for beta-diversity measures and transfer functions using *sed*aDNA datasets. By identifying method combinations that preserve the ecological signal under DNA degradation scenarios, we contribute to the improving of paleoenvironmental reconstructions from *sed*aDNA data.

## Author contributions

TC conceived the study and performed the formal analysis. FK and AL provided empirical datasets and contributed to improve data analysis. TC wrote the paper with input from FK and AL.

## Acknowledgement

TC is supported by the Norwegian Research Council (project number 343086). AL is supported by an assistant research professorship from the IKERBASQUE Foundation.

## Supplementary figures

**Figure S1:**
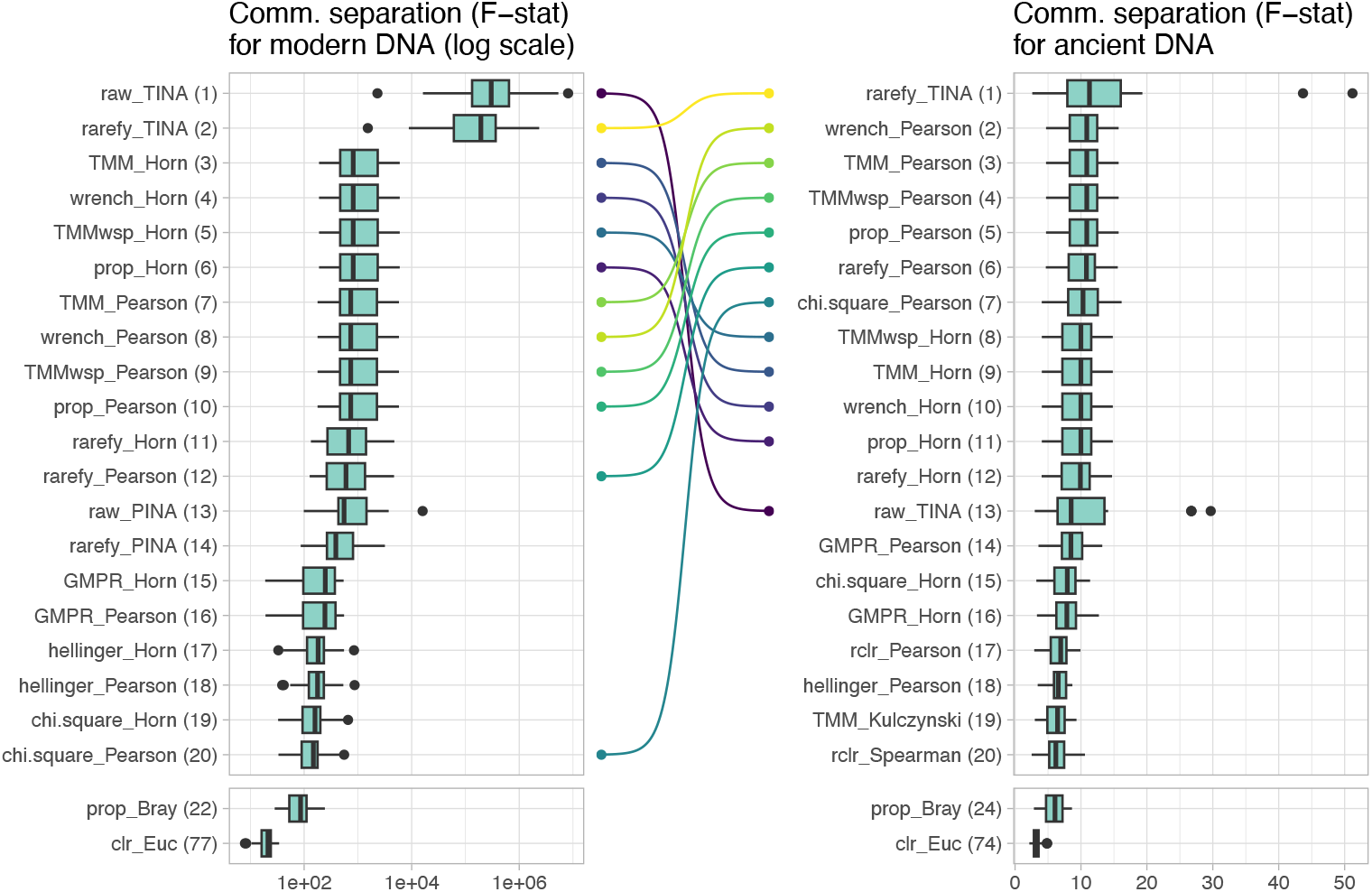
Performance of combinations of DNA read counts transformation and beta-diversity indices for cluster separation in synthetic datasets that contained 2000 taxa. Performance was measured with the F-statistic of PERMANOVA models (higher is better). The left panels show the F-statistic distributions for modern DNA datasets, whereas the right panels show the F-statistic distributions for simulated ancient DNA datasets. Bumps charts connect identical combinations of methods between modern and ancient DNA datasets that are among the 10 best approaches. Numbers between brackets indicate the ranked position of the combinations of count transformation and beta-diversity indices based on the median F-statistic value.

**Figure S2:**
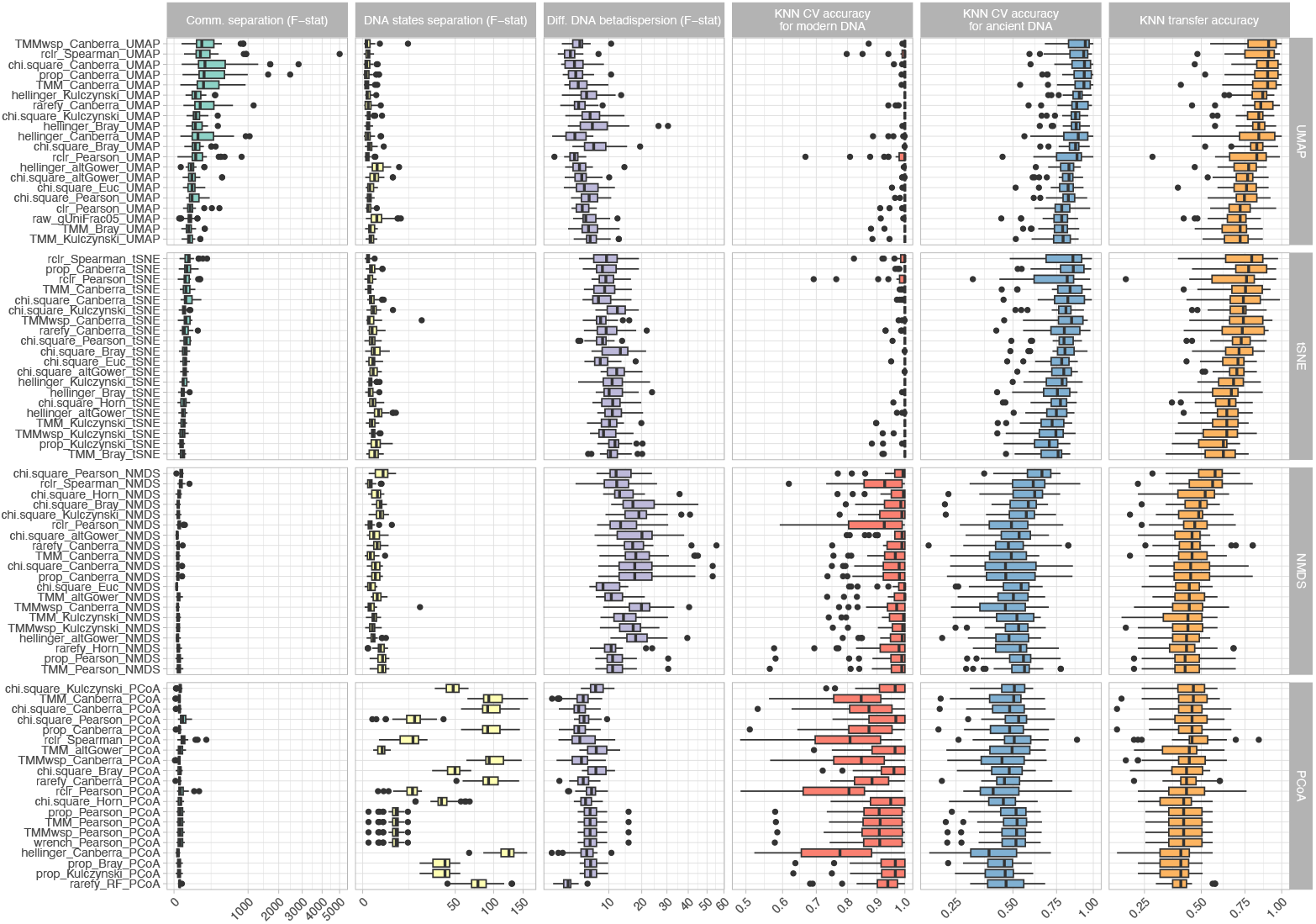
Performance of combinations of DNA read counts transformation, beta-diversity indices and ordinations methods for cluster separation (green panel), DNA status separation (yellow panel), change in betadispersion between DNA state (purple panel), cross validated accuracy on modern datasets (red panel), cross validated accuracy on ancient DNA datasets (blue panel) and transfer function accuracy (orange panel) in synthetic datasets. Only the top 20 combinations that gave best transfer accuracy per ordination method (UMAP, t-SNE, NMDS and PCoA) are shown.

**Figure S3:**
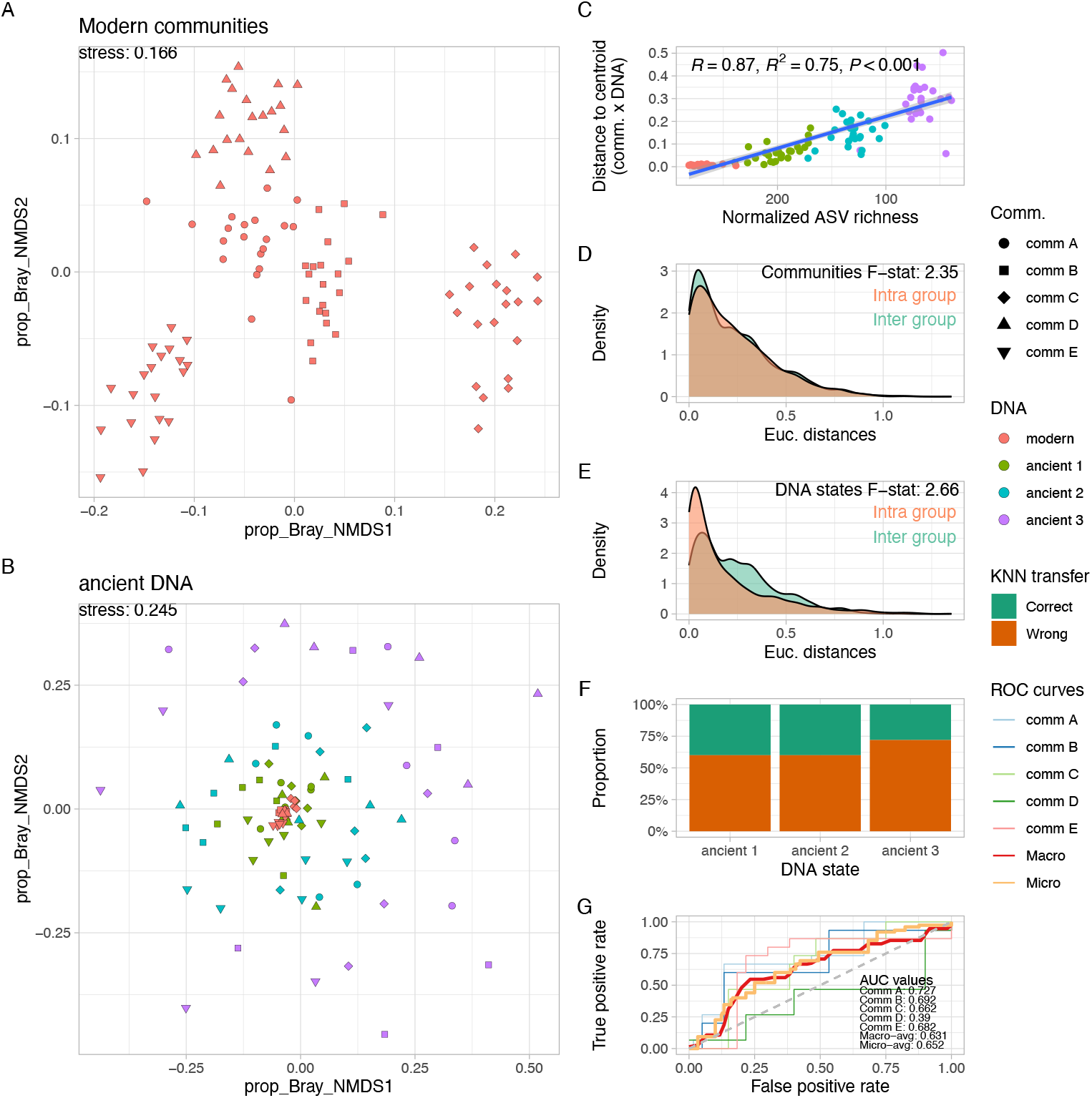
Results obtained on one of the synthetic datasets with five clusters structure with the commonly used proportion transformation technique combined with the Bray-Curtis beta-diversity index and the NMDS ordination method. A) NMDS ordination obtained from the “modern” synthetic dataset. B) NMDS ordination of the same dataset after ancient DNA signal simulation. C) Distances of samples to DNA status centroid (i.e. betadispersion or multivariate variance) as function of normalized taxa (ASV on the axis) richness. D) Distribution of pairwise Bray-Curtis distances within groups (i.e within a given cluster) and between groups (between clusters), as well as the measured F-statistic from a PERMANOVA model. E) Same as D), but here the DNA status was used as factor. F) Proportion of correctly and wrongly classified samples to cluster of origin as a function of ancient DNA effect size. G) Receiver Operating Characteristic (ROC) curves for each individual clusters and overall measure of micro and macro-averages scores.

**Figure S4:**
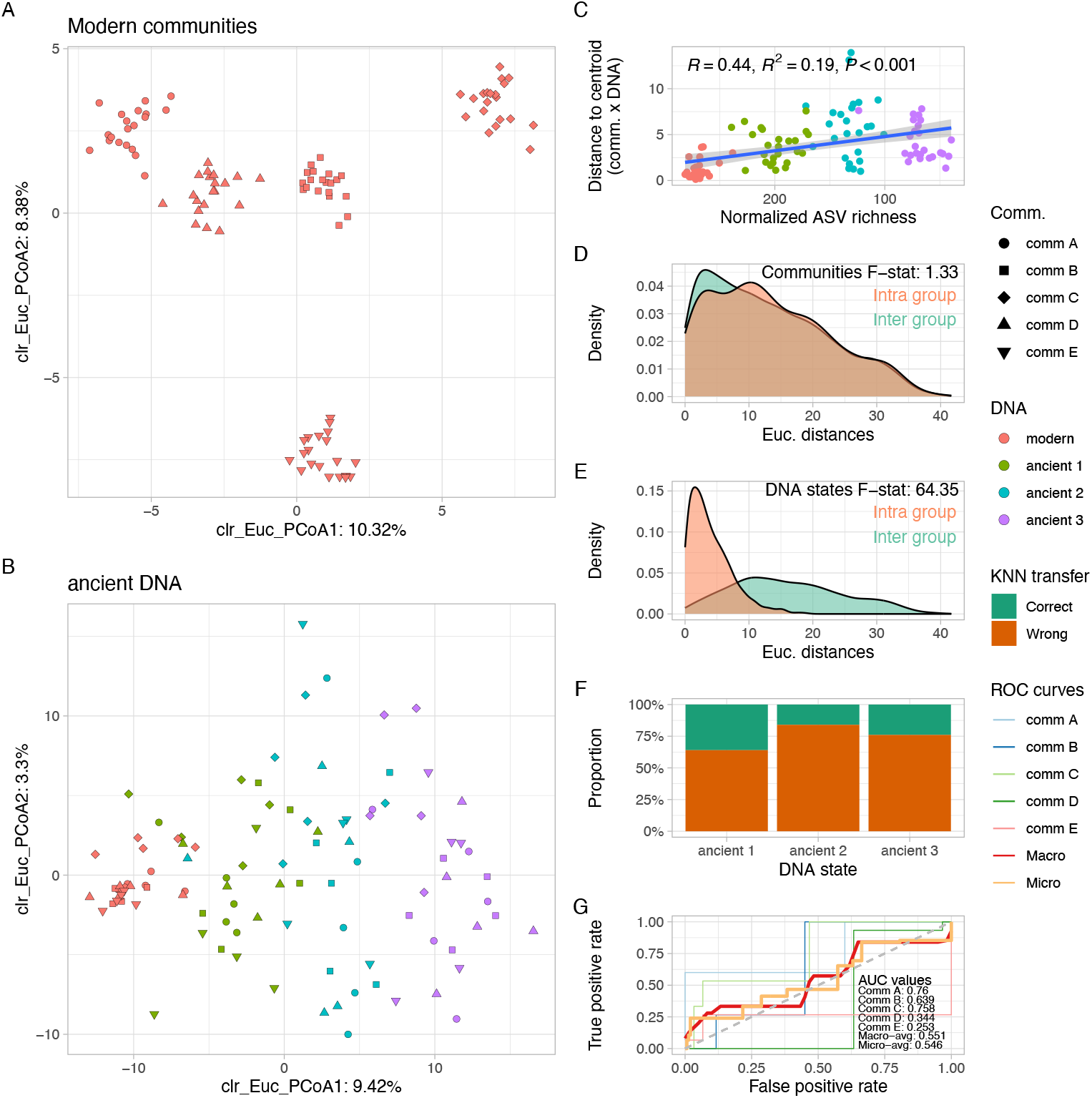
Results obtained on the same synthetic dataset as in Figure S3 with the commonly used center log ratio transformation technique combined with the Euclidean distance metric and the PCoA ordination method. A) PCoA ordination obtained from the “modern” synthetic dataset. B) PCoA ordination of the same dataset after ancient DNA signal simulation. C) Distances of samples to DNA status centroid (i.e. betadispersion or multivariate variance) as function of normalized taxa (ASV on the axis) richness. D) Distribution of pairwise Bray-Curtis distances within groups (i.e within a given cluster) and between groups (between clusters), as well as the measured F-statistic from a PERMANOVA model. E) Same as D), but here the DNA status was used as factor. F) Proportion of correctly and wrongly classified samples to cluster of origin as a function of ancient DNA effect size. G) Receiver Operating Characteristic (ROC) curves for each individual clusters and overall measure of micro and macro-averages scores.

**Figure S5:**
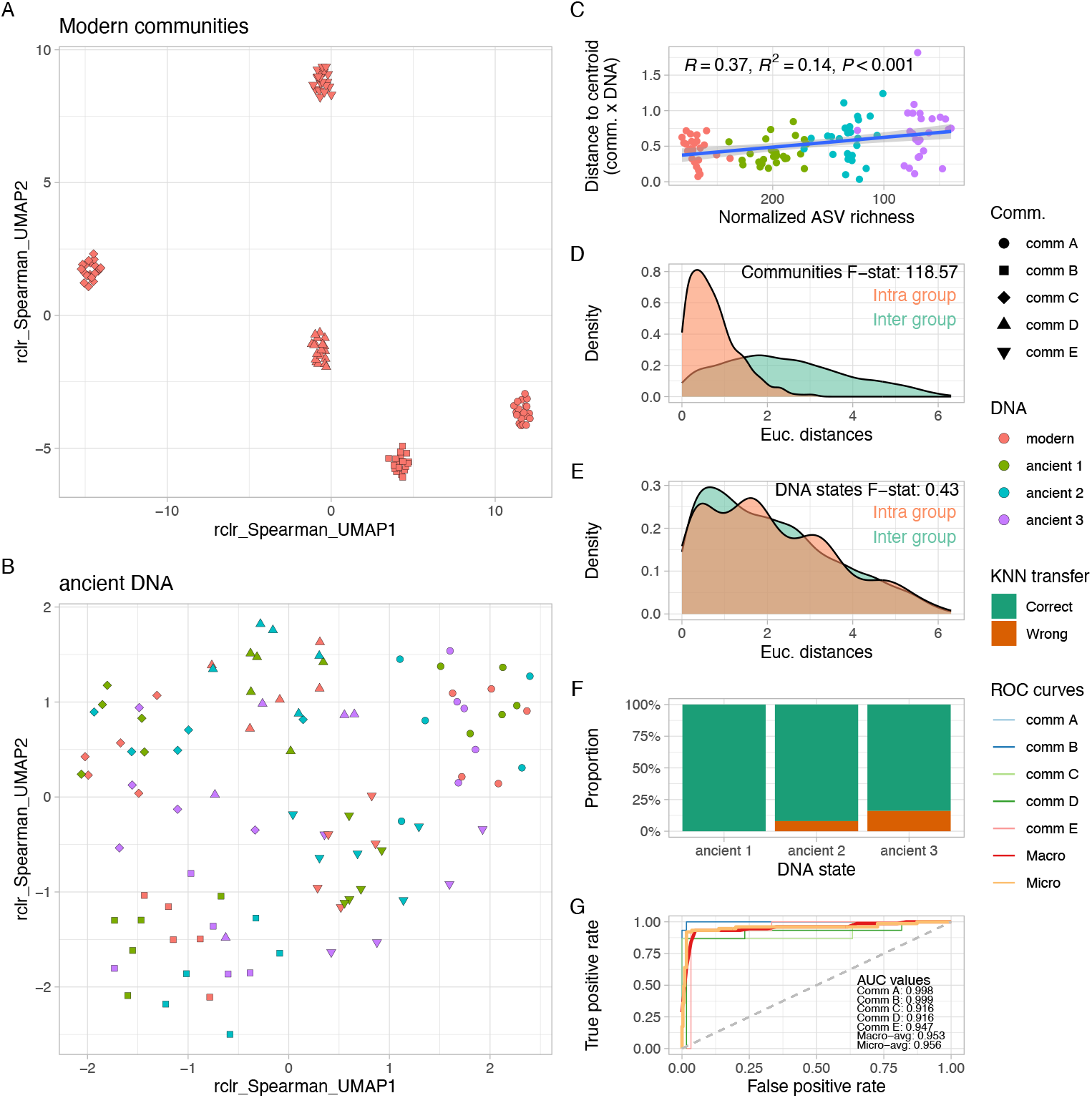
Results obtained on the same synthetic dataset as in Figure S3 with the robust center log ratio transformation technique combined with the Spearman index as a measure of dissimilarity and the UMAP ordination method. A) UMAP ordination obtained from the “modern” synthetic dataset. B) UMAP ordination of the same dataset after ancient DNA signal simulation. C) Distances of samples to DNA status centroid (i.e. betadispersion or multivariate variance) as function of normalized taxa (ASV on the axis) richness. D) Distribution of pairwise Bray-Curtis distances within groups (i.e within a given cluster) and between groups (between clusters), as well as the measured F-statistic from a PERMANOVA model. E) Same as D), but here the DNA status was used as factor. F) Proportion of correctly and wrongly classified samples to cluster of origin as a function of ancient DNA effect size. G) Receiver Operating Characteristic (ROC) curves for each individual clusters and overall measure of micro and macro-averages scores.

## References

1. Guiot J. Transfer functions. IOP Conf Ser Earth Environ Sci 2011;14:10–15. 10.1088/1755-1315/14/1/012008

2. Pflaumann U et al. Glacial North Atlantic: Sea-surface conditions reconstructed by GLAMAP 2000. Paleoceanography 2003;18. 10.1029/2002PA000774

3. Kucera M et al. Reconstruction of sea-surface temperatures from assemblages of planktonic foraminifera: Multi-technique approach based on geographically constrained calibration data sets and its application to glacial Atlantic and Pacific Oceans. Quat Sci Rev 2005;24:951–998. 10.1016/j.quascirev.2004.07.014

4. De Vernal A et al. Dinocyst-based reconstructions of sea ice cover concentration during the Holocene in the Arctic Ocean, the northern North Atlantic Ocean and its adjacent seas. Quat Sci Rev 2013;79:111–121. 10.1016/j.quascirev.2013.07.006

5. Fritz SC et al. Reconstruction of past changes in salinity and climate using a diatom-based transfer function. Nature. 1991. 1991., 352: 706–708

6. Liu B, Cao S. Asynchronous changes in trophic status of a lake and its watershed inferred from sedimentary diatoms of different habitats. Ecol Indic 2018;90:215–225. 10.1016/j.ecolind.2018.03.018

7. Ramstack JM et al. The application of a diatom-based transfer function to evaluate regional water-quality trends in Minnesota since 1970. J Paleolimnol 2003;29:79–94. 10.1023/A:1022869205291

8. Hutchinson GE. Concluding Remarks. Cold Spring Harb Symp Quant Biol 1957;415–427.

9. Lejzerowicz F et al. Ancient DNA complements microfossil record in deep-sea subsurface sediments. Biol Lett 2013;9:20130283. 10.1098/rsbl.2013.0283

10. Epp LS. A global perspective for biodiversity history with ancient environmental DNA. Mol Ecol 2019;28:2456–2458. 10.1111/mec.15118

11. Armbrecht LH. The potential of sedimentary ancient DNA to reconstruct past ocean ecosystems. Oceanography 2020;33:116–123. 10.5670/oceanog.2020.211

12. Capo E et al. Lake Sedimentary DNA Research on Past Terrestrial and Aquatic Biodiversity: Overview and Recommendations. Quaternary 2021;4:6. 10.3390/quat4010006

13. Capo E et al. Is planktonic diversity well recorded in sedimentary DNA? Toward the reconstruction of past protistan diversity. Microb Ecol 2015. 10.1007/s00248-015-0627-2

14. Keck F et al. Assessing the response of micro-eukaryotic diversity to the Great Acceleration using lake sedimentary DNA. Nat Commun 2020;3831. 10.1038/s41467-020-17682-8

15. Monchamp MÈ et al. Paleoecological evidence for a multi-trophic regime shift in a perialpine lake (Lake Joux, Switzerland). Anthropocene 2021;35. 10.1016/j.ancene.2021.100301

16. De Schepper S et al. The potential of sedimentary ancient DNA for reconstructing past sea ice evolution. ISME J 2019;13:2566–2577. 10.1038/s41396-019-0457-1

17. Zimmermann HH et al. Marine ecosystem shifts with deglacial sea-ice loss inferred from ancient DNA shotgun sequencing. Nat Commun 2023. 10.1038/s41467-023-36845-x

18. Armbrecht L et al. Ancient marine sediment DNA reveals diatom transition in Antarctica. Nat Commun 2022;13:5787. 10.1038/s41467-022-33494-4

19. Kjær KH et al. A 2-million-year-old ecosystem in Greenland uncovered by environmental DNA. Nature 2022;612:283–291. 10.1038/s41586-022-05453-y

20. Freeman CL et al. Survival of environmental DNA in sediments: Mineralogic control on DNA taphonomy. Environ DNA 2023. 10.1002/edn3.482

21. Sand KK et al. Importance of eDNA taphonomy and sediment provenance for robust ecological inference: Insights from interfacial geochemistry. Environ DNA 2024;6. 10.1002/edn3.519

22. McMurdie PJ, Holmes S. Waste Not, Want Not: Why Rarefying Microbiome Data Is Inadmissible. PLoS Comput Biol 2014;10. 10.1371/journal.pcbi.1003531

23. Gloor GB et al. It’s all relative: analyzing microbiome data as compositions. Ann Epidemiol 2016;26:322–329. 10.1016/j.annepidem.2016.03.003

24. Quinn TP et al. A field guide for the compositional analysis of any-omics data. Gigascience 2019;8:1–14. 10.1093/gigascience/giz107

25. Kuczynski J et al. Microbial community resemblance methods differ in their ability to detect biologically relevant patterns. Nat Methods 2010;7:813–819. 10.1038/nmeth.1499

26. Bàlint M et al. Millions of reads, thousands of taxa: Microbial community structure and associations analyzed via marker genesa. FEMS Microbiol Rev 2016;40:686–700. 10.1093/femsre/fuw017

27. Martino C et al. A novel sparse compositional technique reveals microbial perturbations. mSystems 2019;4:1–13. 10.1128/msystems.00016-19

28. Kumar MS et al. Analysis and correction of compositional bias in sparse sequencing count data. BMC Genomics 2018;19:799. 10.1186/s12864-018-5160-5

29. Lozupone C et al. UniFrac: An effective distance metric for microbial community comparison. ISME J 2011;5:169–172. 10.1038/ismej.2010.133

30. Schmidt TSB, Rodrigues JFM, Von Mering C. A family of interaction-adjusted indices of community similarity. ISME J 2016;1–17. 10.1038/ismej.2016.139

31. Van Der Maaten LJP, Hinton GE. Visualizing high-dimensional data using t-sne. J Mach Learn Res 2008;9:2579–2605. 10.1007/s10479-011-0841-3

32. McInnes L, Healy J, Melville J. UMAP: Uniform Manifold Approximation and Projection for dimension reduction. 2020.

33. Armstrong G et al. Uniform Manifold Approximation and Projection (UMAP) reveals composite patterns and resolves visualization artifacts in microbiome data. mSystems 2021;6. 10.1128/mSystems.00691-21

34. Milošević D et al. The application of Uniform Manifold Approximation and Projection (UMAP) for unconstrained ordination and classification of biological indicators in aquatic ecology. Sci Total Environ 2022;815. 10.1016/j.scitotenv.2021.152365

35. Dillies MA et al. A comprehensive evaluation of normalization methods for Illumina high-throughput RNA sequencing data analysis. Brief Bioinform 2013;14:671–683. 10.1093/bib/bbs046

36. Weiss SJ et al. Effects of library size variance, sparsity, and compositionality on the analysis of microbiome data. PeerJ Prepr 2015;3:e1408. 10.7287/peerj.preprints.1157v1

37. Pereira MB et al. Comparison of normalization methods for the analysis of metagenomic gene abundance data. BMC Genomics 2018;19:1–17. 10.1186/s12864-018-4637-6

38. Schloss PD. Rarefaction is currently the best approach to control for uneven sequencing effort in amplicon sequence analyses. mSphere 2024;9. 10.1128/msphere.00354-23

39. Zappia L, Phipson B, Oshlack A. Splatter: simulation of single-cell RNA sequencing data. Genome Biol 2017. 10.1186/s13059-017-1305-0

40. Cordier T et al. Patterns of eukaryotic diversity from the surface to the deep-ocean sediment. Sci Adv 2022;8:eabj9309. 10.1126/sciadv.abj9309

41. Oksanen J et al. vegan: Community Ecology Package. 2024. 2024.

42. Cover TM, Hart PE. Nearest Neighbor Pattern Classification. IEEE Trans Inf Theory 1967;13:21–27. 10.1109/TIT.1967.1053964

43. Breiman L. Random Forests. Mach Learn 2001;45:5–32. 10.1023/A:1010933404324

44. Kuhn M. Building Predictive Models in R Using the caret Package. J Stat Softw 2008;28:1–26. 10.1053/j.sodo.2009.03.002

45. Wickham H. ggplot2: Elegant Graphics for Data Analysis. Springer-Verlag New York, 2016.

46. Pedersen TL. patchwork: The Composer of Plots. 2024. 2024.

47. Bray JR, Curtis JT. An Ordination of the Upland Forest Communities of Southern Wisconsin. Ecol Monogr 1957;27:325–349. 10.2307/1942268

48. Aitchison J. The statistical analysis of compositional data. London: Chapman and Hall, 1986.

49. Chari T, Pachter L. The specious art of single-cell genomics. PLoS Comput Biol 2023;19. 10.1371/journal.pcbi.1011288

50. Morton JT et al. Uncovering the horseshoe effect in microbial analyses. mSystems 2017;2:1–7. 10.1128/msystems.00166-16

51. Esling P, Lejzerowicz F, Pawlowski J. Accurate multiplexing and filtering for high-throughput amplicon-sequencing. Nucleic Acids Res 2015;43:2513–2524. 10.1093/nar/gkv107

52. Carøe C, Bohmann K. Tagsteady: A metabarcoding library preparation protocol to avoid false assignment of sequences to samples. Mol Ecol Resour 2020;20:1620–1631. 10.1111/1755-0998.13227

